# The role of transporters and synaptic cleft morphology in glutamate and GABA homeostasis and their effect on neuronal function

**DOI:** 10.1101/670844

**Authors:** Ghanim Ullah

## Abstract

The spatiotemporal dynamics of glutamate and gama-aminobutyric acide (GABA) in the synaptic cleft plays a key role in the signal integration in the brain. Since there is no extracellular metabolism of glutamate and GABA, cellular uptake through transporters and diffusion to extracellular space (ECS) regulates the concentration of both neurotransmitters in the cleft. We use the most up to date information about the transporters and synaptic cleft to model the homeostasis of both glutamate and GABA. We show that the models can be used to investigate the role played by different isoforms of transporters, uptake by different neuronal compartments or glia cells, and key parameters determining the morphology of synaptic cleft in the neurotransmitter concentration in the cleft and ECS, and how they shape synaptic responses through postsynaptic receptors. We demonstrate the utility of our models by application to simple neuronal networks and showing that varying the neurotransmitter uptake capacity and synaptic cleft parameters within experimentally observed range can lead to significant changes in neuronal behavior such as the transition of the network between gamma and beta rhythms. The modular form of the models allows easy extension in the future and integration with other computational models of normal and pathological neuronal functions.

## Introduction

Signal integration in the brain is determined by the amplitude and kinetics of synaptic responses [1, 2], which in turn are controlled by the spatiotemporal dynamics of neurotransmitter concentrations in the synaptic cleft [3, 4, 5]. Among many, glutamate and gamaaminobutyric acide (GABA) are the major excitatory and inhibitory neurotransmitters, and are thereby involved in most aspects of normal brain function including cognition, memory, and learning, and many neuronal disorders [6, 7, 8, 9, 10, 11]. A tight control of both glutamate and GABA in the synaptic cleft and extracellular space (ECS) is therefore crucial for avoiding abnormal neuronal activity [7, 12, 13, 14].

In the absence of extracellular metabolism, cellular uptake maintains low levels of glutamate and GABA concentrations in the cleft and ECS to avoid excitotoxicity or overinhibition in the brain [15, 9, 7, 16, 17]. It has been clear for almost fifty years now that both neurons and astrocytes express different types of transporters to buffer glutamate and GABA from the synaptic cleft and ECS and restore their intracellular pools [7, 18, 10, 14, 16, 19, 20, 21]. It is therefore not surprising that the malfunction of different glutamate and GABA transporters have been linked to several neurological and neuropsychiatric disorders (see for example [7, 10, 16, 22, 23, 24]).

The transporter isoforms and their relative expression levels vary from one cellular compartment to another, from neuron to glia, and between brain regions. For example, out of the five different types of glutamate (excitatory amino acid) transporters (EAATs), EAAT2 is responsible for about 95% of the total glutamate uptake and is expressed mainly in astrocytes and axonal terminals at 10:1 ratio [19]. Similarly, the densities of EAAT1 and EAAT2 in cerebellum are more than double and 10% of those in the CA1 region of the hippocampus respectively [19]. EAAT3 is predominantly expressed in dendrites only while EAAT4 and EAAT5 are found only in Purkinje cells and retina respectively and do not play a major role in glutamate uptake in other brain regions [7, 18].

Another key factor in shaping neurotransmitter concentrations and hence synaptic currents is the geometry of synaptic cleft. For example, variations of the glutamate concentration in the cleft and differences in the potency of vesicles released from different locations on the active zone are shown to be the two main contributors to the quantal variability at single glutamatergic synapse [25]. Modulation of glutamate mobility in the cleft and spillover to the ECS has also been shown to shape *α*-amino-3-hydroxy-5-methyl-4-isoxazolepropionic acid (AMPA)-induced synaptic currents [26]. Similarly, the height of the synaptic cleft has been correlated with the optimal amplitude of the synaptic currents [5]. The morphology of glia surrounding the synapse also seems to play a key role in the spatiotemporal profile of glutamate concentration in the cleft and spillover to the ECS [27].

To summarize, the vast variability in the distribution and function of transporters and parameters shaping the morphology of synaptic cleft play crucial roles in the spatiotemporal dynamics of glutamate and GABA, and are therefore key to our understanding of how they affect synaptic currents and neuronal function. However, current experimental techniques are too limited to investigate all these variables one by one or simultaneously, and warrant biophysical computational models - the subject of this paper.

## Methods

We use two separate models that describe the electrical properties and neurotransmitter dynamics of a glutamatergic and a GABAergic neuron.

### Membrane Model

The equations describing the membrane potential of each cell are described in section “Membrane Model” of Supplementary Information S1 Text.

To assess the interplay of membrane potential and neurotransmitter dynamics, we add a range of glutamate and GABA–related processes to the model. There are significant simplifications at work and we like to emphasize that the goal is to present a general framework for the role played by key electrochemical and morphological variables in the glutamate and GABA homeostasis.

### Glutamate–Related Processes

Glutamate is a neurotransmitter that is released into the cleft of a synaptic connection when the presynaptic, i.e. signal–sending, neuron depolarizes. Glutamate binds to the NMDA and AMPA receptors of the postsynaptic neuron and can thereby initiate an action potential. After binding to a receptor the transmitter is free again and can bind another time or diffuse into the ECS. Neurons and glia (astrocyte) cells clear glutamate by taking it up from the cleft or from the ECS. For an overview of glutamate–related processes we refer the reader to [28, 7, 29, 18]. The basic formalism for glutamate homeostasis presented in this section is modified and extended from our previous work [30].

#### Glutamate Release and Diffusion

During an action potential, about 3,000 glutamate molecules are released into the synaptic cleft [31, 32, 33, 29]. Glutamate release also depends on the remaining glutamate, 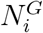, in the presynaptic terminals. We remark that it must be carried in vesicles (3,000 glutamate molecules per single vesicle) to be released properly. Initially, the releasable amount of glutamate will be at the maximal level, 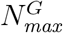, but with high frequency spiking it is reduced [34]. Thus the amount of glutamate released per action potential at a single synapse is given by

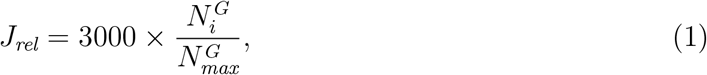

where 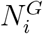 is the amount of glutamate in the neuron that is available for release and is given by the difference between 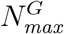 and the total glutamate released, 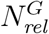.

Neurons and astrocytes take up the released glutamate from the cleft and ECS, but at first the buffered glutamate is not enclosed in vesicles—it cannot be used for synaptic signals right away. Buffered glutamate gets recycled to produce new vesicles. The intracellular diffusion and recycling of the buffered glutamate into new vesicles slowly recover 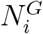.

Glutamate is released into the synaptic cleft that is located at the dendritic terminal (see Fig. 1). Its size is given by its height *h* and radius *r*. We assume a half–spherical shape and obtain the following cleft volume *ω*_*c*_ (Fig. 1 inset):

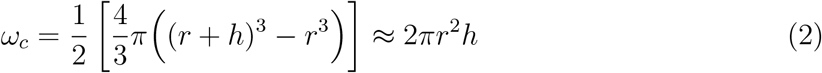

**Figure 1:**
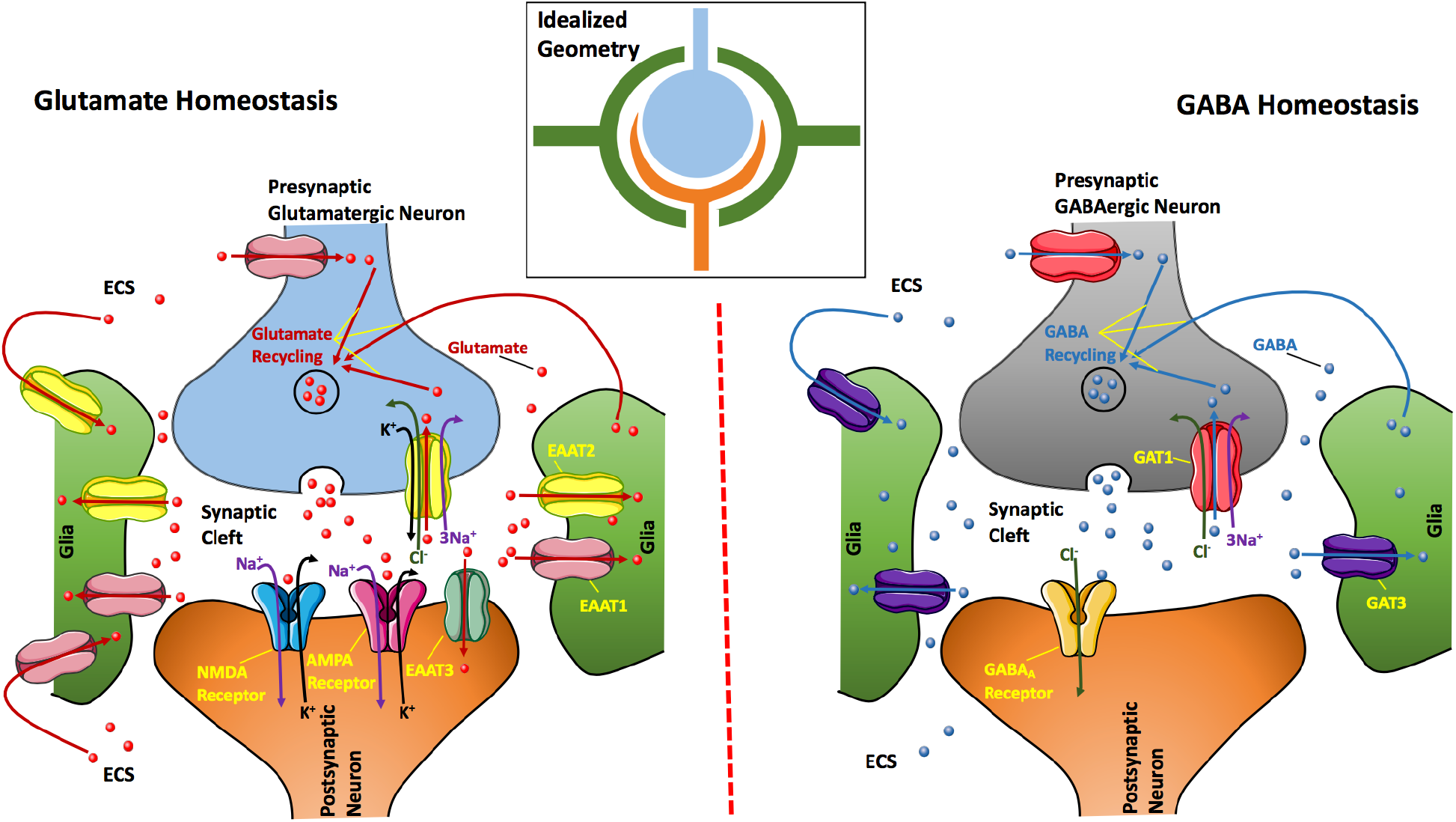
Schematic of the key processes used in the model for glutamate (left) and gaba (right) homeostasis. When a signal travels from the presynaptic glutamatergic(gabaergic) neuron along the axon to a postsynaptic neuron, glutamate(gaba) is released at the axonal terminal (bouton). The terminal is separated from the neighboring neuron by the synaptic cleft. Glutamate(gaba) can excite(inhibit) postsynaptic neuron by binding to NMDA/AMPA receptors to bring in Na^+^ and release K^+^ (GABA_A_ receptor to bring in Cl^−^) on the spine located at the dendrite of the receiving neuron. At the same time the presynaptic neuron can receive glutamate(GABA)–mediated signals from other neurons through its own dendritic terminals. Glutamate(GABA) is buffered from the cleft by the bouton of the presynaptic neuron through EAAT2(GAT1)and glia surrounding the synaptic cleft through EAAT1 and EAAT2 (GAT3). The spine of the postsynaptic neuron also buffers glutamate from the cleft through EAAT3. Glutamate(GABA) can also diffuses to the ECS where it is buffered by neuron through EAAT1(GAT1) and glia through EAAT1 and EAAT2(GAT3). EAAT1, EAAT2, and EAAT3 transport one glutamate molecule along with three Na^+^ and one Cl^−^ into the cell while releasing one K^+^. GAT1 and GAT3 transport one GABA molecule along with three Na^+^ and one Cl^−^ into the cell. The inset on the top shows idealized geometry of the synapse considered in the model.

Typical values for *r*, *h*, and other morphological parameters are given in Table B of Supplementary Information S1 Text. Terms of the order 𝒪 (*h*^2^) and higher are omitted, because *h* ≪ *r*.

Astrocytes reach out to the dendritic cleft creating the so–called glial envelope [35] (Fig. 1). We estimate the whole volume *ω*_*en*_ that is enclosed in this envelope to be three times as large as the cleft (Fig. 1 inset):

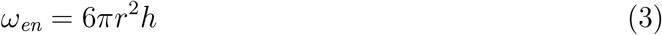

In the following, we will assume that neurotransmitter in the cleft spreads into the whole envelope immediately after its release, i.e., the concentrations in the cleft and the envelope are the same and we refer to them synonymously. With this assumption we will only distinguish between glutamate concentrations in the ECS and the cleft and denote them by *G*_*e*_ and *G*_*c*_, respectively. The release of ∆*N*^*G*^ glutamate molecules leads to a cleft concentration of

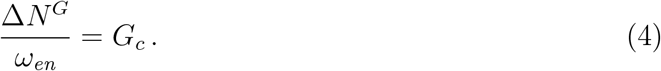

If we assume a baseline level of nearly zero, 3,000 molecules increase the concentration by 1.3 mM (considering *r* = 110nm and *h* = 20nm [36, 37, 5]).

One important mechanism that clears glutamate from the cleft is diffusion. To estimate the glutamate diffusion rate we note that the cross section area, *A*_*σ*_, for fluxes from the envelope into the ECS is only 5% of the outer spherical surface of the dendritic connection, because 95% are covered by the glial envelope [35]:

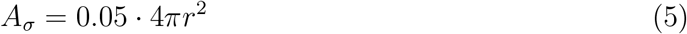

Let *D*_*G*_ be the glutamate diffusion coefficient [38] and ∆*x* the cutoff distance from the synapse at which *G*_*e*_ is in a steady state [35]. Then we get the following flux of glutamate out of the cleft:

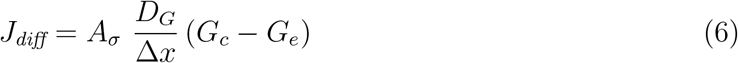

#### Glutamate Uptake

Binding to AMPA and NMDA receptors is not a clearance mechanism and glutamate will be re–released into the cleft. What clears glutamate is uptake by the neuron and astrocyte. The mathematical description of cellular glutamate uptake is formulated in terms of the density of available binding sites on the neuron or astrocyte cell *B*. As a chemical reaction scheme, glutamate uptake can be pictured as follows [35, 39]:

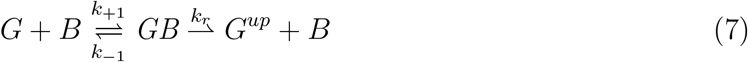

We apply this scheme with different rates to model five uptake scenarios: uptake from the cleft into axonal terminals (boutons), astrocyte, and dendritic spines, and from the ECS into the neuron and astrocyte. *G* is the glutamate concentration in the cleft or the ECS, and *B* is the neural or astrocytic concentration of free binding sites through which the neurotransmitter can be transported into the cells. Bound glutamate is denoted by *GB*. It can either be re–released or taken into the cell. Both these processes leave a free binding site. Buffered glutamate is denoted by *G*^*up*^. Under the assumption that this reaction chain is stationary with a constant transporter concentration, *B*, the following uptake velocity, *v*_*e*/*c*→*cell*_, of glutamate from ECS/celft into the cell, i.e. the velocity of the process *G* ⇀ *G*^*up*^ can be derived (see [40] for this kind of derivation):

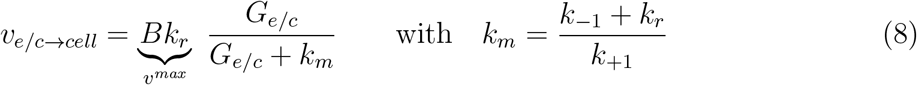

The velocity is measured in [femtomole/ms], and we note that in this scheme the uptake capacity is described by spatial density *B* of binding sites, which is proportional to the surface density *ρ*^*B*^ of binding sites in the cellular membrane. With a cycling rate (*CycRate*) of 30 molecules/s for glutamate transporters [41, 42, 43], *v*^*max*^ for a given uptake area (*A*^*up*^) becomes,

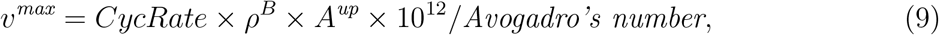

where 10^12^/*Avogadro’s number* converts the number of molecules/s to femtomole/ms.

There are five different types of glutamate transporters: EAAT1, EAAT2, EAAT3, EAAT4, and EAAT5. In this study we will restrict our modeling to glutamate uptake in hippocampus. Out of the five transporters, EAAT4 and EAAT5 are expressed in Purkinje cells and retina respectively, and do not play a major role in glutamate uptake in hippocampus [7]. EAAT1 is expressed in astrocytes surrounding the synapses with surface density 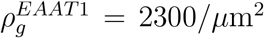 in hippocampus [19, 18]. EAAT2 represents about 95% of the total glutamate activity in the adult brain [14, 24] and is expressed both in astrocytes and axonal terminals (boutons) at density 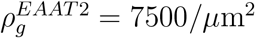 and 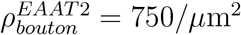 respectively [18]. EAAT3 is predominantly expressed in spines and dendrites with density 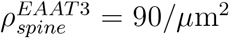[18]. Thus we substitute *ρ*^*B*^ in Eq. (9) by 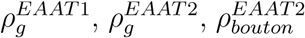, and 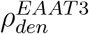 to model the uptake by astrocyte through EAAT1, astrocyte through EAAT2, bouton through EAAT2, and spine through EAAT3 from synaptic cleft respectively. The observed affinity (*k*_*m*_) for EAAT1, EAAT2, and EAAT3 are 7 – 20*µ*M, 12 – 18*µ*M, and 8 – 30*µ*M respectively [18]. We use 13.5*µ*M, 15*µ*M, and 19*µ*M (the mean) as the affinity for EAAT1, EAAT2, and EAAT3 respectively.

As mentioned above, glutamate uptake also depends on the surface area *A*^*up*^ that is available for uptake. For astrocyte uptake from the envelope we note that astrocyte cells cover the synapse from the outside which is approximately one spherical surface area (see Fig. 1). Neural uptake is through those parts of the neural membranes facing this astrocyte envelope (one spherical surface), and from below and above the cleft (two half–spherical surfaces). This makes the uptake area for bouton and spine one spherical surface each with radius *r*. That is, 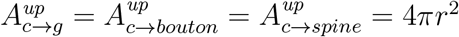.

To model the uptake from ECS by astrocyte through EAAT1 and EAAT2, we simply replace *A*^*up*^ by the total membrane area of the astrocyte 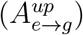 in Eq. (9). Similarly, the neuronal uptake from ECS is modeled by using the total membrane area of the neuron 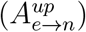. Since, EAAT2 is mostly expressed in terminals and dendrites make most of the surface area of the neuron, we only include the uptake by neuron through EAAT3 [18]. The values of and logic behind the morphological parameters are given in Table B and section “Morphology” of Supplementary Information S1 Text respectively.

Altogether glutamate dynamics are described by eight dynamical variables: the average amount in the cleft 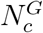, the total amount in the ECS 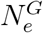, the total glutamate released 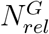 (glutamate that is not available for release), the net uptake from all synaptic clefts (multiple synapses are involved in a single action potential) by astrocyte 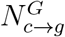, bouton 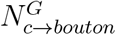, and spine 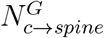, and the net uptake from ECS by astrocyte 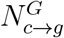 and neuron 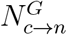. The rate equations are

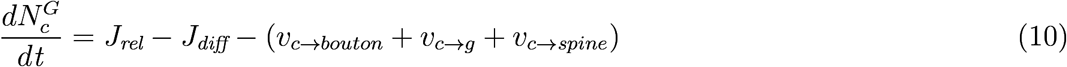

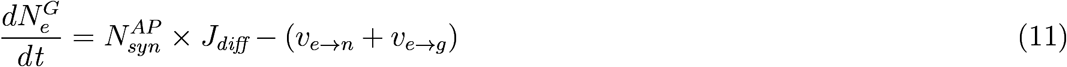

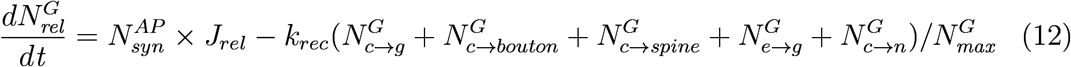

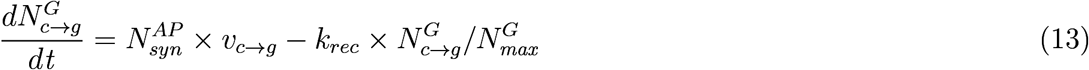

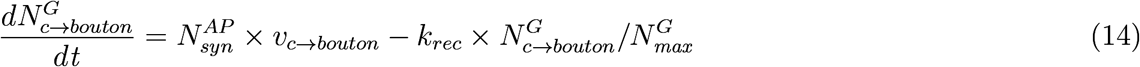

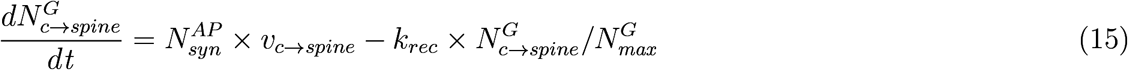

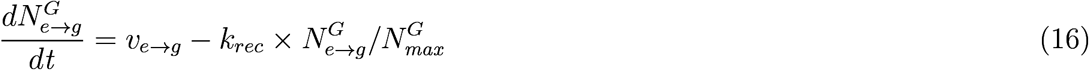

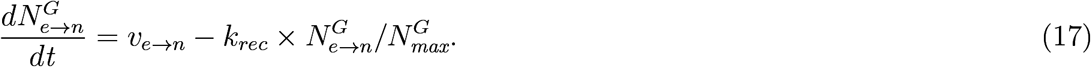

Where 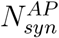 is the number of synapses involved in a normal action potential and *k*_*rec*_ is the recycle rate for glutamate. Stevens and Tetsuhiro [34] estimated that in hippocampal neurons, on average a single synapse has approximately 20 releasable sites that take about 10 s to replenish. Considering 10,000 synapses per neuron, this will lead to an overall recycle rate *k*_*rec*_ of 9.9635 × 10^−5^femtomole/ms. The amount of recycled glutamate is proportional to the relative uptake by the compartment.

If an estimate for the amount of glutamate uptake by individual compartments is not desired, Eqs. (13–17) can be combined into one rate equation modeling the total buffered glutamate 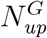, that is

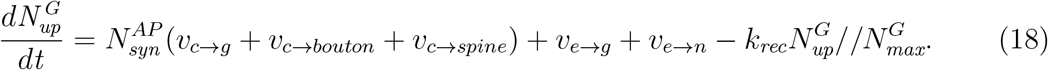

The expression 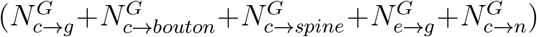 in Eq. (12) changes to 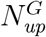 and the number of dynamical variables reduces to four. Various parameters used in the glutamate model and relevant receptors (see next section) are given in Table C of Supplementary Information S1 Text.

#### NMDA and AMPA Receptor Binding

Glutamate at high concentrations will bind to the receptors on the postsynaptic dendrites. Specifically, it excites a neuron by binding to the NMDA and AMPA receptors. Computational models for the effect of receptor gates on action potentials have been developed for normal action potential events involving approximately 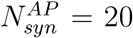 synapses. The effective receptor conductance in such scenario is estimated to be in the range 1 × 10^−8^ − 6 × 10^−7^ mS for NMDA 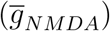 and 3.5 × 10^−7^ − 1 × 10^−6^ mS for 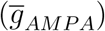 receptors [44]. Since in this paper we consider normal spiking, we keep 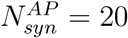 and use 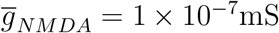 and 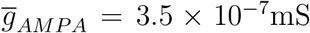. Both values are divided by the total cell membrane area to obtain the values in mS/cm^2^. In more excited states such as epileptic seizure and spreading depolarization where the tissue is flooded with glutamate, the number of synapses involved and the total maximum conductances for both receptors will be significantly larger.

NMDA and AMPA receptor gates open for Na^+^ and K^+^ ions, and the opening probability of the particular gate is described by the gating variables *r*_*NMDA*_ and *r*_*AMPA*_. Their dynamics are given by a Hodgkin–Huxley–like formalism with an additional dependence on *G*_*c*_[44]:

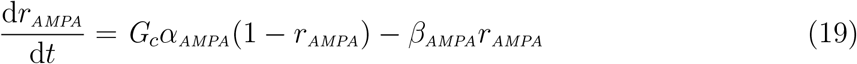

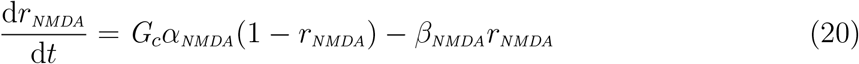

The parameters *α*_*NMDA/AMPA*_ and *β*_*NMDA/AMPA*_ were estimated by Destexhe et al. [44] by fitting both models to experimental data. When compared to detailed Markov chain models with several gating states and taking into account the desensitization of the receptor, these simpler models were shown to fit the observed postsynaptic currents through NMDA and AMPA receptors equally well [44]. The receptor currents are given as

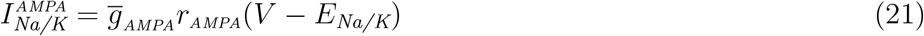

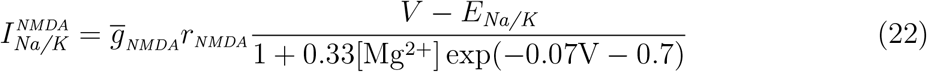

Uptake of glutamate goes along with ion cotransport [28]. Thus the rate equations for membrane potential change accordingly. For the neuron, one molecule of glutamate is accompanied by three Na^+^ and one Cl^−^, while it releases one K^+^. These contributions can be converted to the cotransport currents

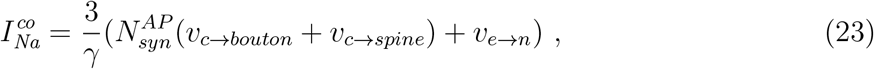

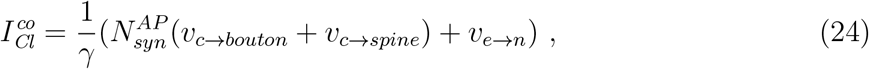

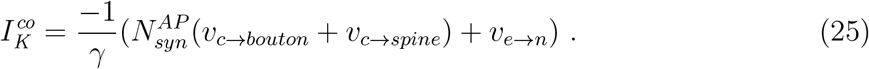

Where *γ* = *A*_*m*_/*F* changes ion flux (femtomole/ms) to current density (*µA*/*cm*^2^). 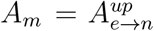 and *F* is the membrane area of the neuron and Faraday’s constant respectively. These and the AMPA and NMDA currents must be added to the rate equation for the membrane potential, i.e. we need to replace

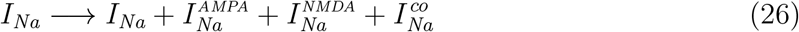

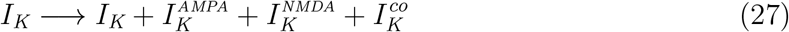

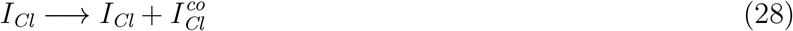

in Eq. (1S) in Supplementary Information Text 1S.

### GABA–Related Processes

Our formalism for GABA homeostasis is similar to glutamate with some key differences outlined below.

#### GABA Release and Diffusion

We assume that the number of GABA molecules released at a single synapse during an action potential and the maximum amount of releasable GABA is similar to glutamate. We also assume that the morphological parameters describing the synaptic cleft, neuronal, astrocytic, and extracellular compartments remain the same. However, the diffusion coefficient of GABA in the cleft is 0.51*µm*^2^/*ms*, almost double that of glutamate [45].

#### GABA Uptake

The equation for the uptake velocity of GABA from ECS/cleft to neuron or astrocyte is also similar to Eq. (8). That is,

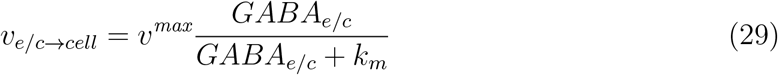

However, the values of *v*^*max*^ and *k*_*m*_ are different. The observed turnover rate (*CycRate* in Eq. 9) for GABA transporter 1 (GAT1) is 79 – 93*s*^−1^ [46]. We use the mean of this range as *CycRate* for GABA transporters.

The three main transporters involved in GABA uptake are GAT1, GAT2, and GAT3. It is generally believed that GAT1 is the most important of the three transporters and is primarily present on GABAergic neurons but also to some extent on astrocytes [20]. Thus GABAergic activity is mainly terminated by GABA transport into the presynaptic terminals [47, 48]. GAT2 is expressed neonatally in the brain but its expression decreases with time and are mostly confined to the meninges [14, 17, 16, 49] and will be ignored in our formalism. In contrast to GAT1, GAT3 is considered to be selectively expressed in astrocytes throughout the brain [17, 50].

Chiu et al. [51] showed that the presynaptic boutons of GABAergic interneurons in the hippocampus has a surface density of of GAT1 molecules, 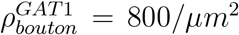. Given that we do not have any significant evidence of GAT molecules on the spines, we consider GABA uptake by spines to be zero, i.e. 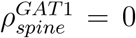 [52, 16]. However, considering 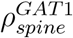 to be up to 20% of 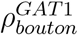 does not change the time-course of GABA concentration in the cleft significantly. Although the density of GAT3 has not been measured to our knowledge, only 10-20% GABA is taken up by astrocytes [17, 53]. Thus, we take the density of GAT3 in astrocytic membrane to be 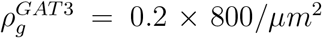 for both astrocytic uptake from cleft and ECS (but different uptake ares 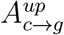 versus 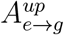, see Eq. 9). We also use an indirect approach for estimating GAT1 density on neuronal membrane other than boutons in hippocampus since a direct measurement is not available. In cerebellum, Chiu et al. [51] estimated GAT1 density to be 1340/*µm*^2^ and 677*/µm*^2^ in boutons and axons respectively, giving a ratio of 0.5052. A similar ratio was found in cortical slices as well. We assume this to be a rough estimate of GAT1 density in neuronal membrane. Thus, we take 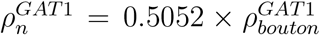 with uptake area 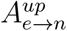. However, this ratio can be varied in the model to see how it affects the overall behavior of GABA homeostasis. The observed affinity (*k*_*m*_) for GAT1 and GAT3 is 7*µ*M and 0.8*µ*M respectively [14, 54, 55, 56, 10].

All the above considerations result in a set of rate equations for GABA homeostasis similar to Eqs. (10–17). With the assumption *v*_*c→spine*_ = 0, the rate equation for 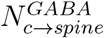 (similar to Eq. 15) is not required. Like glutamate homeostasis, the detailed model for GABA with seven dynamical variables reduces to a four-variable model if the overall GABA uptake is desired instead of the uptake by individual compartments.

Various parameters used in the GABA model and relevant receptor (see next section) are given in Table D of Supplementary Information Text 1S.

#### GABA_A_ Receptor Binding

GABA binds to GABA receptors on the postsynaptic neuron. Specifically, it inhibits a neuron by binding to the ionotropic receptor GABA_A_ or metabotropic receptor GABA_B_. In this study, we only consider GABA_A_ receptors and adopt the model developed by Destexhe et al. [44]. GABA_A_ receptor opens for Cl^−^ ions with the opening probability of the gate described by the gating variable 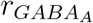. The dynamics of the gate is given by a Hodgkin–Huxley–like formalism with an additional dependence on the GABA concentration *GABA*_*c*_ in the cleft [44]:

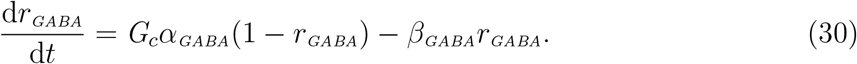

The receptor current is given as

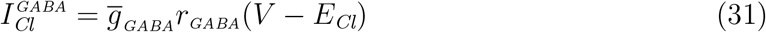

One molecule of GABA is accompanied by three Na^+^ and one Cl^−^ [57, 58], leading to the GABA cotransport currents

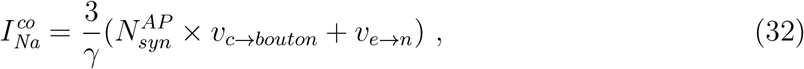

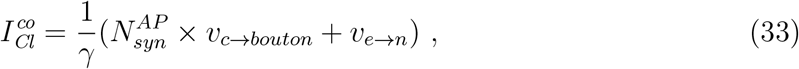

These and the GABA_A_ current must be incorporated in the rate equation for the membrane potential by replacing

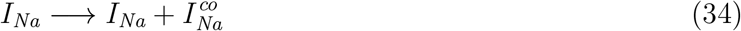

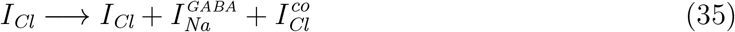

in Eq. (1S) in Supplementary Information Text 1S.

#### Numerical Methods

The model equations for both glutamate and GABA homeostasis were implemented in Fortran 90/95. To facilitate the dissemination of these results, codes reproducing the key results in this paper will be provided as Supplementary Information S2 Text upon the acceptance of the paper for publication and will also be archived at modelDB [59].

## Results

### Glutamate and GABA homeostasis as functions of transporters density

The above equations allow us to investigate the role played by various uptake sources and the key parameters characterizing the morphology of synaptic cleft in the neurotransmitter dynamics and how they modulate synaptic currents and neuronal function. Fig. 2 shows the evolution of glutamate and GABA concentrations in synaptic cleft and ECS, and different uptake mechanisms. In the model, neurotransmitter is released in quantum of 3,000 molecules as the neuronal membrane potential crosses a fixed threshold of 0 mV during the rising phase of the action potential (not shown) onset. Glutamate (Fig. 2A) and GABA (Fig. 2C) concentration in the cleft peaks at a fixed value of about 1.3 mM, irrespective of the density of the transporters. The peak value in the ECS on the other hand, decreases as we double the density of transporters. The time to decay for neurotransmitter concentration both in the cleft and ECS also decreases. In the absence of transporters, the only way for concentration in the cleft to decay is to diffuse to the ECS. Thus *G*_*e*_ and *GABA*_*e*_ first increase and then plateau as there is no uptake from ECS.

**Figure 2:**
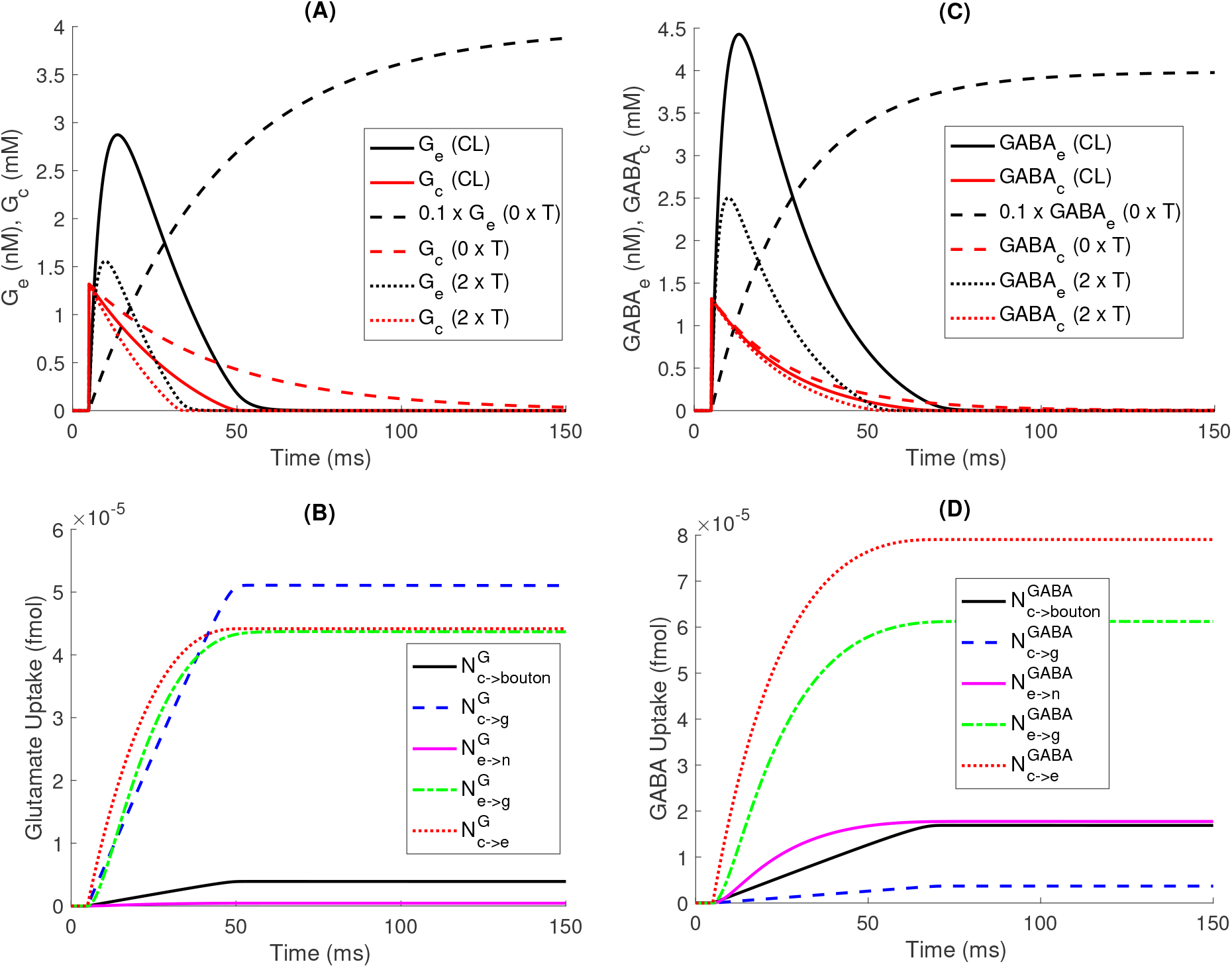
Glutamate and GABA homeostasis at normal transport (CL), no transport (0×*T*), and double that of normal transport (2 × *T*). Glutamate (A) and GABA concentrations (C) in the ECS and cleft. Note that *G*_*e*_ (A) and *GABA*_*e*_ (C) at no transport are multiplied by 0.1 for better visualization. Glutamate (B) and GABA (D) uptake into various compartments from cleft and ECS under the three conditions.

While similar in many ways, there are obvious differences in the glutamate and GABA homeostasis. There is a significantly larger spillover of synaptic GABA to the ECS as compared to glutamate (compare the peak concentrations in ECS in Fig. 2A & C). In case of glutamate, the dominant mechanisms for uptake from the cleft are the astrocyte and diffusion to the ECS (Fig. 2B). In case of GABA on the other hand, diffusion to ECS predominantly controls the uptake from the cleft Fig. 2D. That is why, the decay time for *G*_*c*_ shows a stronger dependence on transporter density when compared to *GABA*_*e*_. In both cases, astrocyte is the main sink for uptake from ECS. Uptake from the cleft by the spine of postsynaptic neuron is very small and is not shown.

The change in the temporal dynamics of *G*_*c*_ has serious consequences for both *I*_*AMPA*_ and *I*_*NMDA*_. In Fig. 3, we show the currents through the receptors and transporters from the simulations in Fig. 2. While the peak value of *I*_*AMPA*_ does not change, its duration increases significantly as we reduce the transporter density (Fig. 3A). Both the peak value and duration of *I*_*NMDA*_ change when we change the uptake capacity. Whereas there seems to be a slight increase in the peak value, the duration of the current through GABA receptor increases significantly as we decrease the density of GAT1 and GAT3 (Fig. 3C). Glutamate (Fig. 3B) and GABA (Fig. 3D) transporters also result in a small net inward neuronal current. Although small (e.g. 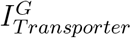 is two orders of magnitude smaller than *I*_*NMDA*_) on the single action potential scale, these currents could play a significant role in neuronal function in highly excited states such as seizure and spreading depolarization [30].

**Figure 3:**
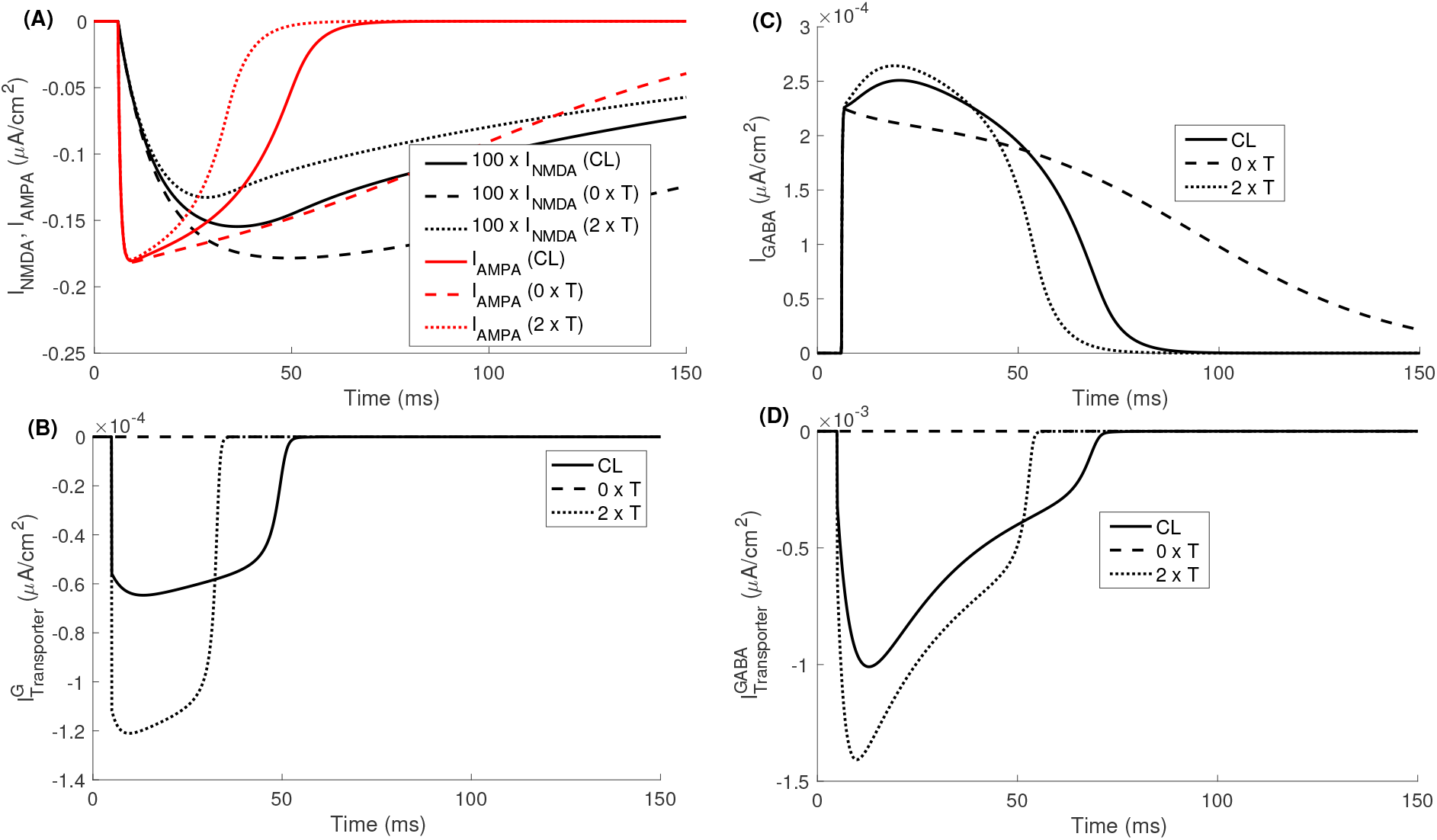
Neuronal currents through receptors and transporters at normal transport (CL), no transport (0 × *T*), and double that of normal transport (2 × *T*). Currents through NMDA and AMPA receptors (A), total net neuronal current through glutamate (EAAT1, EAAT2, and EAAT3) transporters (B), current through GABA_A_ receptor (C), and GABA transporter (GAT1) (D) under the three conditions. Note that *I*_*NMDA*_ (A) is augmented 100 times for better visualization.

To have a more quantitative look, we repeat the simulation in Figs. 2 & 3 by changing the transporters density in small increments and show how the peak values and duration for which the neurotransmitter concentrations and synaptic currents change. The amplitude of *G*_*e*_ (Fig. 4A), *I*_*NMDA*_ (Fig. 4C, top panel), *I*_*AMPA*_ (Fig. 4C, bottom panel), and *GABA*_*e*_ (Fig. 4E) decreases exponentially as we increase transporters density. A similar behavior is shown by the duration (time for a variable to drop from its peak value to less than 5% of the peak value) of *G*_*c*_, *G*_*e*_ ((Fig. 4B), *I*_*NMDA*_ (Fig. 4D, top), *I*_*AMPA*_ (Fig. 4D, bottom), *GABA*_*c*_, *GABA*_*e*_ (Fig. 4F, main), and *I*_*GABA*_ (Fig. 4F, inset). Interestingly, there is a slight decrease in the amplitude of *I*_*GABA*_ (Fig. 4F, inset). The peak values of *G*_*c*_ and *GABA*_*c*_ do not change significantly and are not shown.

**Figure 4:**
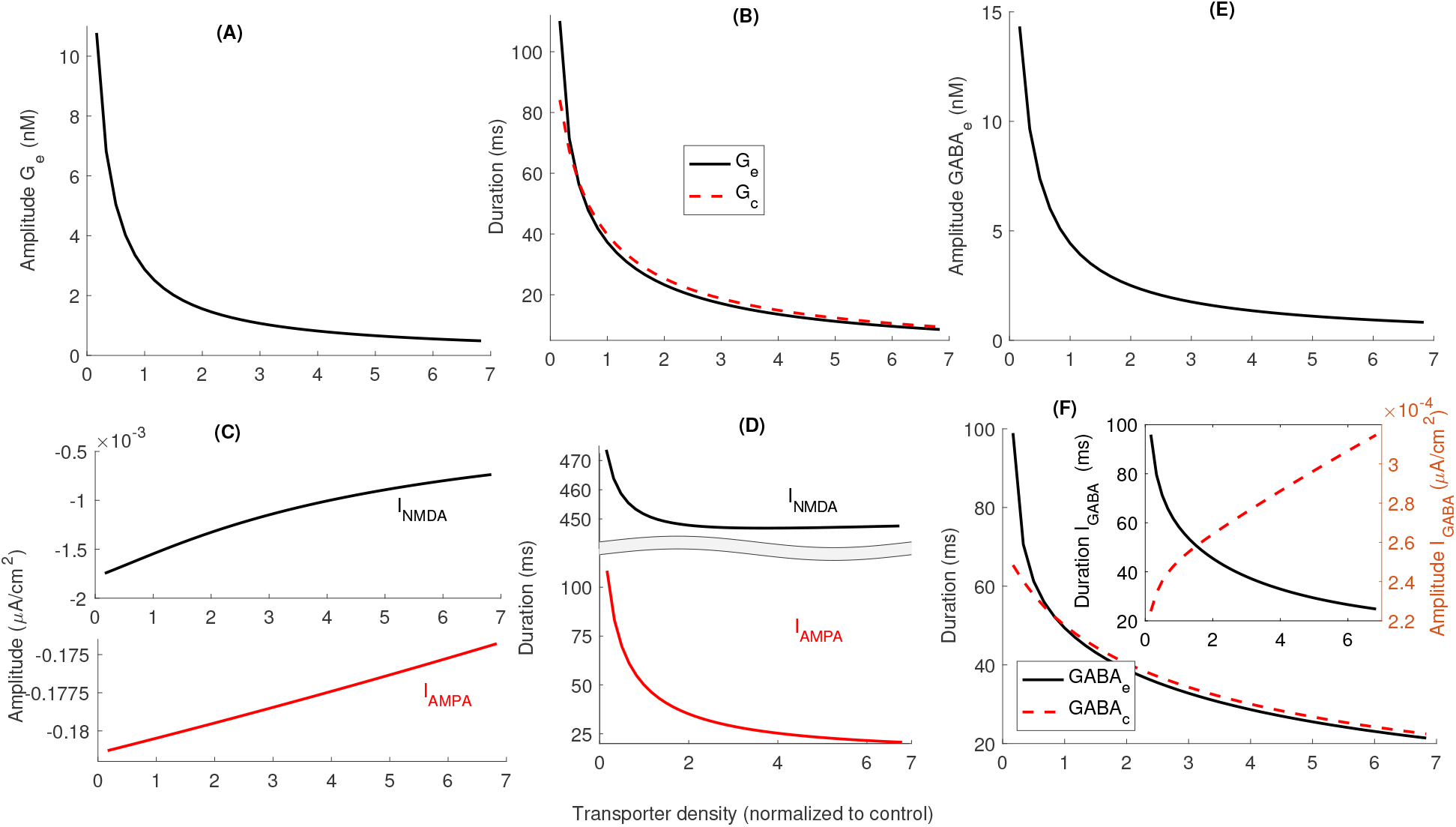
Neurotransmitter homeostases and synaptic currents as functions of transporter density. The amplitude of *G*_*e*_ (A) and duration of *G*_*e*_ and *G*_*c*_ (B) decreases with increasing density of glutamate transporters, resulting in the smaller amplitude of current through NMDA (C, top) and AMPA (C, bottom) receptors. (D) Duration of the current through NMDA (top) and AMPA (bottom) also decreases as we increase transporters density. (E, F main) Same as (A, B) but for GABA homeostasis. (F, inset) Duration decreases (solid line, left axis) while amplitude increases slightly (dashed line, right axis) as we increase transporter density. Note that a more positive value of peak *I*_*NMDA*_ and *I*_*AMPA*_ means smaller current since the inward current due to positive charge is taken as negative in our convention.

### Glutamate and GABA homeostasis as functions of synaptic cleft morphology

In addition to the density of transporters, the size and morphology of the synaptic cleft play a crucial role in the spatiotemporal dynamics of neurotransmitters and related currents. In the above results, we assumed that only 5% of the outer spherical surface of the dendritic connection is available for neurotransmitter diffusion from the cleft to ECS. The remaining area is covered by glial envelope. By incrementally increasing this cross-sectional area from 1 to 100% of the the outer spherical surface of the dendritic connection, we notice that the peak value of *G*_*e*_ (Fig. 5A) and *GABA*_*e*_ (Fig. 5E) increases while that of *I*_*NMDA*_ and *I*_*AMPA*_ (Fig. 5C) decreases. Although the change is very small, the peak value of *I*_*GABA*_ first increases and then decreases (Fig. 5F, inset). The duration of neurotransmitter concentrations in the ECS and cleft (Fig. 5B & F, main), and the synaptic currents (Fig. 5D, & F inset) decreases exponentially as we increase *A*_*σ*_. Note that the change in the dynamics of *I*_*GABA*_ is dominated by the change in duration and the slight increase in the amplitude at lower *A*_*σ*_ values does not make a huge difference.

**Figure 5:**
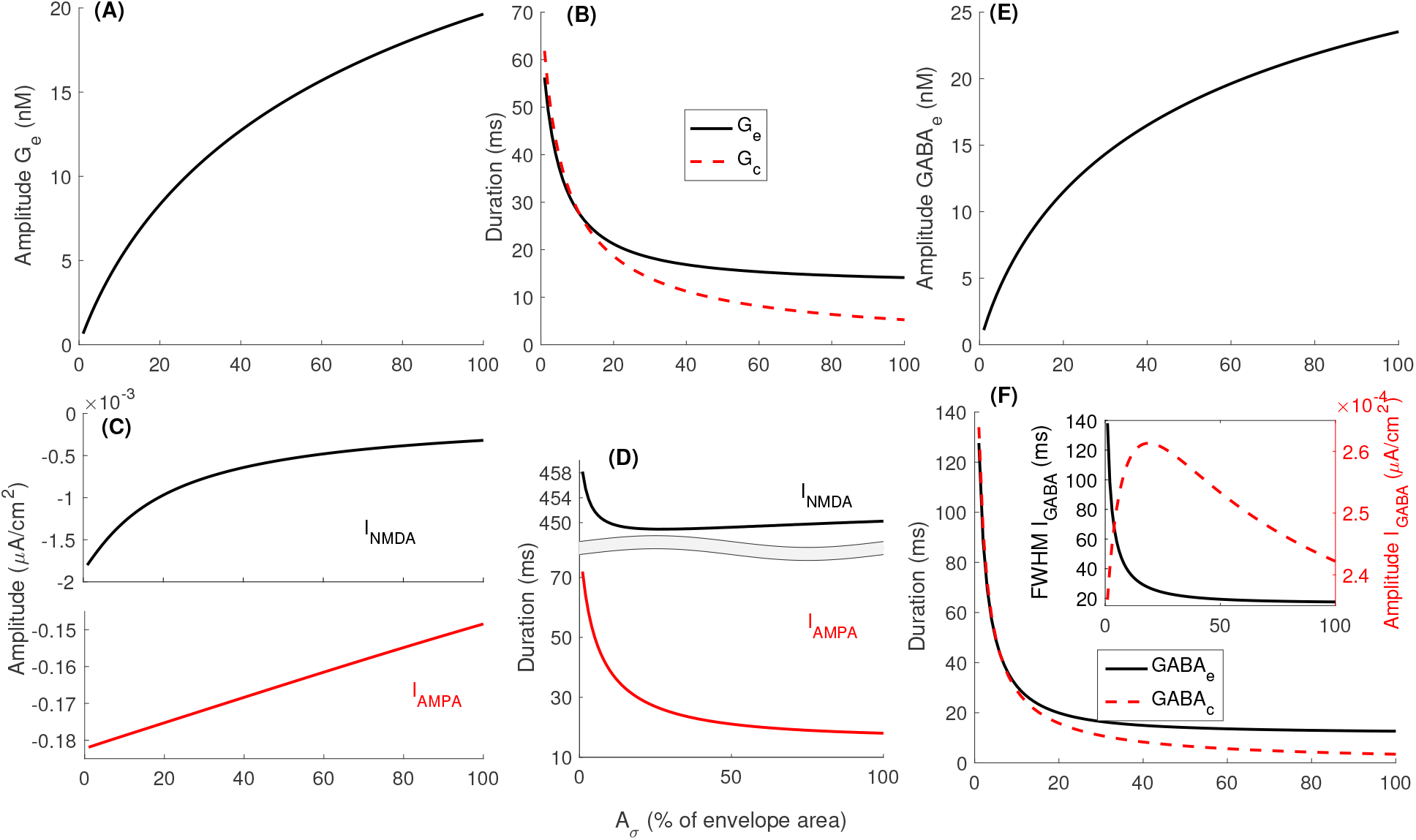
Dependence of neurotransmitter homeostases and synaptic currents on *A*_*σ*_. Peak *G*_*e*_ (A), duration of *G*_*e*_ and *G*_*C*_ (B), amplitude of *I*_*NMDA*_ (top) and *I*_*AMPA*_ (bottom) (C), and duration of *I*_*NMDA*_ (top) and *I*_*AMPA*_ (bottom) (D). (E, F main) Same as (A, B) but for GABA homeostasis. (F, inset) The duration (solid line, left axis) of *I*_*GABA*_ decreases while the amplitude first increases and then decreases slightly (dashed line, right axis) as we increase *A*_*σ*_. Note that a value of peak *I*_*NMDA*_ and *I*_*AMPA*_ closer to zero means smaller current.

The radius of the synapse (radius of the bouton, spine, and glial envelope) and the height of the cleft are the other key parameters characterizing the morphology of the cleft. Both these parameters vary significantly from one synapse to another. For example, the bouton diameter in cortical and hippocampal glutamatergic [60, 61, 62, 63] and GABAergic [64] neurons vary from 200 nm to more than 1.5*µ*m. Although not as dramatic as the bouton size, considerable variability in the cleft height ranging from 10 to 35 nm in different brain regions has also been observed [65, 5, 66, 67, 68]. As shown in Fig. 6, the peak value of glutamate and GABA concentrations both in ECS (Fig. 6A & I) and cleft (Fig. 6C & K) decreases while the duration increases (Fig. 6B, D, J, & L)) as we increase the hight of the cleft. A similar behavior is shown by *I*_*NMDA*_ (Fig. 6E & F), *I*_*AMPA*_ (Fig. 6G & H), and *I*_*GABA*_ (Fig. 6M & N). Both the amplitude and duration of all these variables decrease as we increase the radius of the synapse (horizontal axes). Note that increasing the radius of the synapse affects neurotransmitter in two ways; (1) the number of transporters increases (we keep the density of transporters fixed), leading to lower concentration, and (2) the cleft volume increases, leading to lower concentration for the same number of neurotransmitter molecules. We remark that our model does not incorporate the potential increase in the electrical resistance of the intra-cleft medium as we decrease *r* or *h* that could counter the affect of reducing the cleft size. As was shown by Savtchenko and Rusakov [5], the interplay between these two opposing mechanisms would lead to an optimal height of the synaptic cleft.

**Figure 6:**
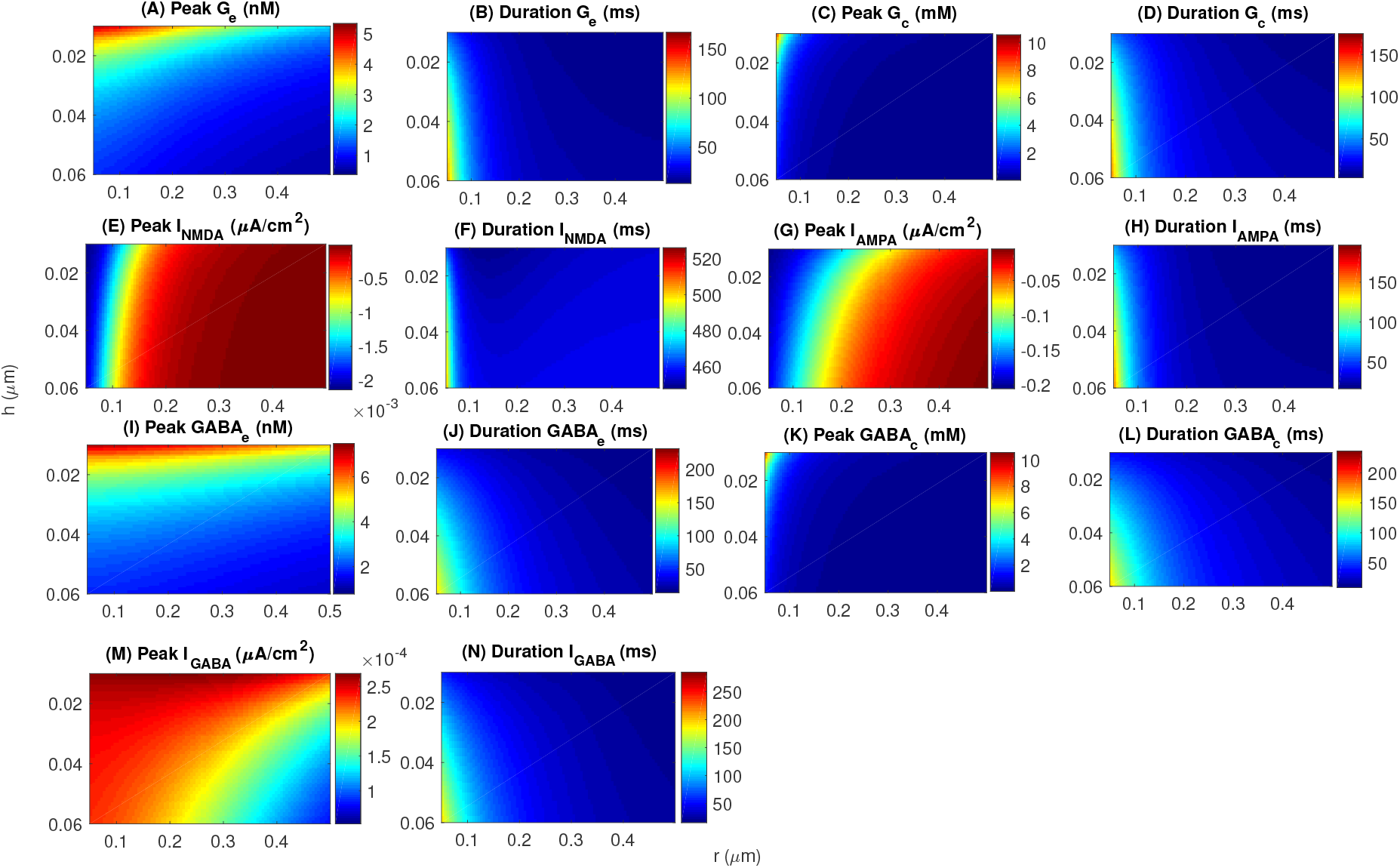
Glutamate, GABA homeostases, and related synaptic currents as functions of the bouton radius (*r*) and height of the synaptic cleft (*h*). Note that a peak value of *I*_*NMDA*_ and *I*_*AMPA*_ closer to zero means smaller current.

### The effect of uptake and synaptic morphology on neuronal spiking

Next, we investigate how the four key parameters discussed in the previous section (transporter density, *A*_*σ*_, *r*, and *h*) affect the excitability of the neuron by modeling two separate small networks of synaptically coupled neurons using the membrane model discussed in the “Methods” section. The first network consists of two glutamatergic neurons while the second network includes two GABAergic neurons. Both neurons in each system are synaptically coupled to each other and receive a unifrom stimulus current. The frequency of one neuron from each network is shown in Fig. 7. The frequency of excitatory postsynaptic neuron decreases from 34 *s*^−1^ to 26 *s*^−1^ as we increase *A*_*σ*_ or transporter density (Fig. 7A). An opposite effect is seen on the function of inhibitory neuron where the frequency increases from almost 13 *s*^−1^ to 26 *s*^−1^ (Fig. 7B). While the effect of cleft height is mild, changing the synaptic radius significantly affects the spiking frequency. The frequency of glutamatergic neuron varies in the range of 34 *s*^−1^ to 26 *s*^−1^ (Fig. 7C) while that of GABAergic neuron varies from less than 15 *s*^−1^ to more than 25 *s*^−1^ (Fig. 7D) as we vary these two parameters.

**Figure 7:**
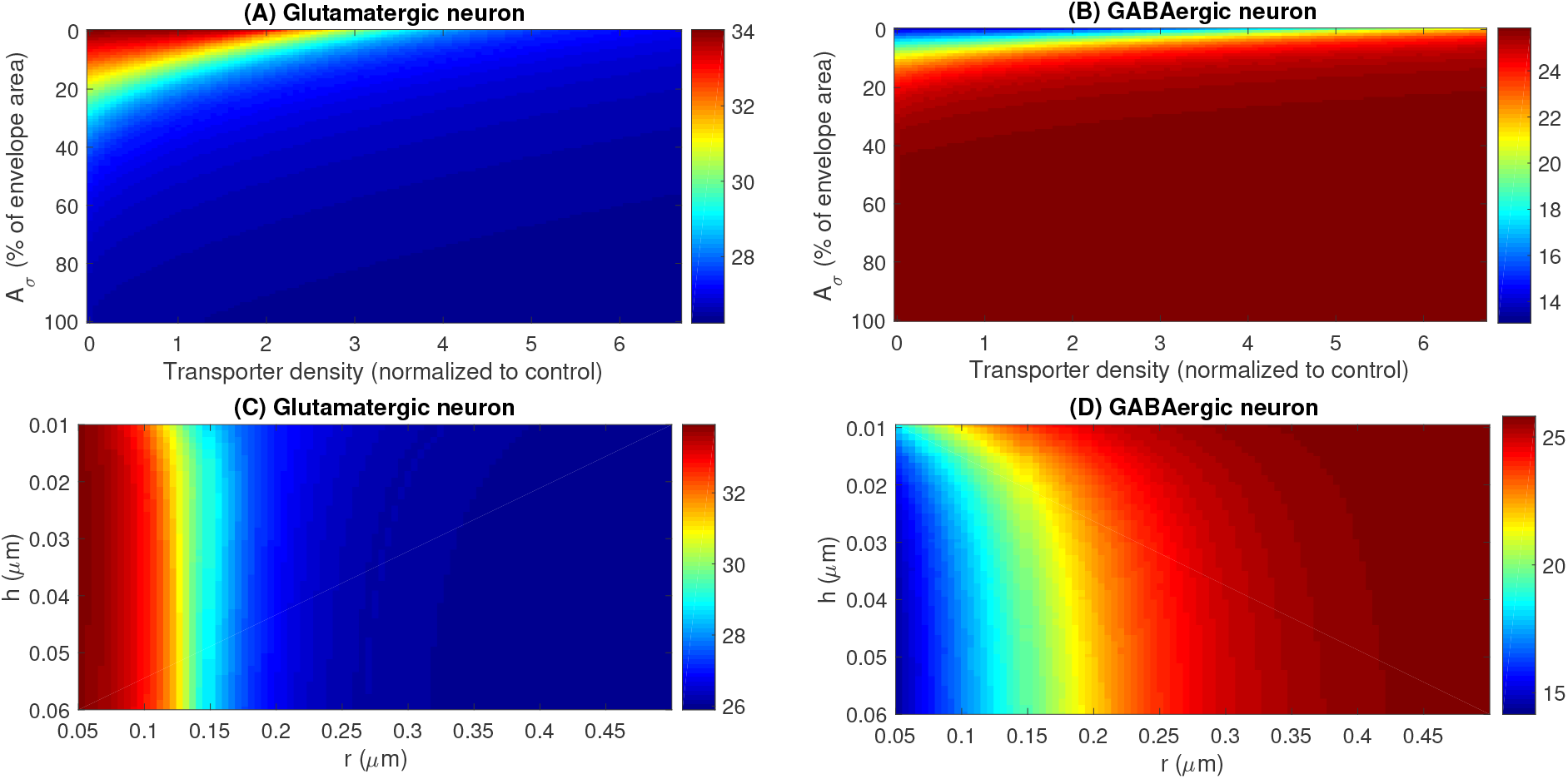
The effect of transporter density, *A*_*σ*_, bouton radius (*r*), and height of the synaptic cleft (*h*) on the spiking frequency of a small excitatory (consisting of two glutamatergic neurons) and a small inhibitory (consisting of two GABAergic neurons) network. Each neuron is stimulated using a current *I*_*stim*_ = 1.75*µA*/*cm*^2^.

## Discussion

Four key considerations were made while developing the models presented above. First, an effort was made to integrate most up to date information about the transporters into the models. For example, the values for affinities, densities, spatial, and cellular distributions of various transporters, the main parameters shaping the synaptic clefts, and the equations modeling the kinetics of various receptors are all based on experimental observations. Second, the model should enable us to delineate the roles of different transporter isoforms, cellular compartments, cell types, and key morphological parameters in neurotransmitter concentration and synaptic/non-synaptic currents. Third, the model equations are formulated so that integration with HH-type formalism for normal neuronal function and/or models investigating the dynamics of ion concentrations in various conditions etc is relatively straight forward. For example, our neurotransmitter models can be easily incorporated in network models studying neuronal rhythms or models for ion concentration dynamics and cell swelling in pathological conditions such seizure and spreading depolarization during migraine, stroke, and traumatic brain injury etc [69, 30, 70, 71]. The models can also be easily used for other brain regions and extended to other types of synapses. The fourth key goal was to leave room for future extension of the models. For example, the recycling of neurotransmitter is modeled by a single term that ignores the complex glutamate-glutamine cycle. The modular form allows easy coupling of our equations to models for glutamate-glutamine cycle or the role of TCA cycle-related variables in neurotransmission [72]. Similarly, our equations can also be coupled to models for neurochemical mechanisms in neurons to investigate the effect of glutamate decarboxylase (GAD) inhibition or vesicular transporters down-regulation on the brain function [73]. GAD inhibition has been shown to cause convulsions and susceptibility to induced seizures [74, 75, 76]. Vesicular GABA transporters knockout also leads to spontaneous and induced seizures [77, 78], and are being considered as potential treatment targets for temporal lobe epilepsy [79].

We use a simple example to demonstrated the utility of our models and show that varying the uptake capacity either by changing the density of transporters or cycling rate can change the frequency of a simple glutamatergic neuronal network from less than 30 *s*^−1^ to well over 30 *s*^−1^. In terms of brain rhythms, this change is significant enough to cause the transition of the network from gamma to beta rhythm and vice versa [80, 81]. A similar effect can be seen when the parameters controlling the morphology of synaptic cleft are varied. These parameters have an opposite effect on a network of GABAergic neurons when compared to glutamatergic network. Given that different brain rhythms are shaped by the interplay between excitatory and inhibitory neuronal networks [82], the uptake mechanism and synaptic cleft morphology could potentially have a more prominent role in shaping the activity of a network consisting of both glutamatergic and GABAergic neurons. Furthermore, the effect in highly excited states such as seizure and spreading depolarization will be more dramatic. We remark that the extent by which the uptake capacity and synaptic cleft parameters are varied in our simulations are well in the experimentally observed range. This clearly highlights the importance of incorporating the neurotransmitter homeostases in neuronal models.

## Acknowledgements

This work was supported by NIH grant R01 AG053988 awarded to GU.

## Supporting Information Legends

**S1 Text.** Neuronal membrane potential model and model parameters.

**S2 Text.** The codes reproducing the key results in this paper will be provided after the acceptance of the manuscript for publication.

